# Using approximate Bayesian computation to quantify cell-cell adhesion parameters in a cell migratory process

**DOI:** 10.1101/068791

**Authors:** Robert J. H. Ross, R. E. Baker, Andrew Parker, M. J. Ford, R. L. Mort, C. A. Yates

## Abstract

In this work we implement approximate Bayesian computational methods to improve the design of a wound-healing assay used to quantify cell-cell interactions. This is important as cell-cell interactions, such as adhesion and repulsion, have been shown to play an important role in cell migration. Initially, we demonstrate with a model of an *ideal* experiment that we are able to identify model parameters for agent motility and adhesion, given we choose appropriate summary statistics. Following this, we replace our model of an ideal experiment with a model representative of a practically realisable experiment. We demonstrate that, given the current (and commonly used) experimental set-up, model parameters cannot be accurately identified using approximate Bayesian computation methods. We compare new experimental designs through simulation, and show more accurate identification of model parameters is possible by expanding the size of the domain upon which the experiment is performed, as opposed to increasing the number of experimental repeats. The results presented in this work therefore describe time *and* cost-saving alterations for a commonly performed experiment for identifying cell motility parameters. Moreover, the results presented in this work will be of interest to those concerned with performing experiments that allow for the accurate identification of parameters governing cell migratory processes, especially cell migratory processes in which cell-cell adhesion or repulsion are known to play a significant role.

## 1 Introduction

Cell-cell interactions are known to play an important role in several cell migration processes. For example, multiple different cell-cell interactions, such as cell-cell signalling and cell-cell adhesion [1], have been identified as promoting metastasis in breast cancer. Repulsive interactions mediated via ephrins on the surface of neural crest stem cells are known to coordinate the early stages of melanoblast migration away from the neural tube [2]. More fundamentally, it is hypothesised that the emergence of cell-cell interactions over one billion years ago helped establish the necessary conditions for multicellular organisms [3].

A well-established approach for studying cell migration is to construct an individual-based model (IBM) to simulate the cell migratory process of interest [4–8]. Typically, this involves using a computational model to simulate a population of agents on a two-dimensional surface, or in a three-dimensional volume. The agents in the IBM represent cells, and each agent is able to move and interact with other agents in the IBM. In this work we use an IBM to simulate a wound-healing assay^1^, an experiment commonly used for studying cell motility [9–11].

If an IBM is an *effective*^2^ representation of a cell migration process it can be used for a number of purposes. One such purpose for an IBM is to perform in *silico* experiments to test scientific hypotheses. For instance, a recent study used an IBM to demonstrate that a simple mechanism of undirected cell movement and proliferation could account for neural crest stem cell colonisation of the developing epidermis in the embryonic mouse [4]. Other studies involving IBMs have tested hypotheses concerning the influence of matrix stiffness and matrix architecture on cell migration [12], and the mechanism by which cranial neural crest stem cells become ‘leaders’ or ‘followers’ in the embryonic chick to allow their collective migration [6–8].

IBMs can also be used to *identify* parameters in experimental data (with the caveat that the parameters are model-dependent). The reasoning behind using an IBM to identify parameters in experimental data is as follows: if an IBM is an effective representation of an experiment, then the parameter values the IBM requires to reproduce the experimental data may be representative of the parameter values in the biological process that is the focus of the experiment^3^. For instance, the value of a parameter that describes cell proliferation rate. Even if the parameter values in the parameterised IBM are not representative of the parameter values in the biological process, the parameterised IBM may still be used to make predictions about the process of interest by performing *in silico* experiments, as described above. These predictions can then be experimentally tested.

Alternatively, if the IBM is an effective representation of an experiment (i.e. the experimental data can be reproduced), but the parameters of the IBM are not identifiable, this may suggest the experiment is not well-designed (that is, if the experiment has been designed to estimate parameters). By parameters not being identifiable it is meant that different parameter values in the IBM can reproduce the same experimental data. If this is the case, the IBM can then be used to suggest improvements to the experiment’s design, namely by altering the IBM design such that the IBM parameters become identifiable. These alterations can then be applied to the experiment to improve parameter identifiability. For example, a recent study using an IBM has examined the time-points at which data should be collected from an experiment to maximise the identifiability of IBM parameters [11]. Other theoretical work has shown how to maximise the information content of an experiment by choosing an appropriate experimental design [13].

The focus of our study is to determine the experimental conditions, and experimental data, required for the accurate identification of cell motility and adhesion parameters in a wound-healing assay. To do so we employ approximate Bayesian computation (ABC), a probabilistic approach whereby a probability distribution for the parameter(s) of interest is generated, as opposed to a point estimate [10, 14, 15]]. Although ABC is well-established in some fields, for instance in population genetics [16], its applicability for IBMs representing cell migration is still an area of active research [10, 11]. Recent studies combining ABC and IBMs have been able to identify motility and proliferation rates in cell migratory processes [10], and improve the experimental design of scratch assays [11]. However, as far as we are aware nobody has used ABC methods to examine the experimental conditions, and experimental data, required for the accurate identification of cell motility and adhesion parameters in a wound-healing assay.

Other methods to identify parameters in experimental data using IBMs also exist. For instance, a standard approach is to generate point estimates of model parameters that best reproduce statistics of the experimental data in the IBM. For example, the generation of motility and proliferation rates for agents in an IBM representing a biological process [4]. This approach, while applicable in some circumstances, often gives no insight into how much uncertainty exists in the parameters chosen, a factor that can be of importance when analysing biological systems. For example, relationships between parameter uncertainty and system robustness are thought to be connected in biological function at a systems level [17].

The outline of this work is as follows: in Section 2 we introduce the IBM and define the cell-cell interactions we implement. We also outline the method of ABC, and the summary statistics we use to analyse the IBM output. In Section 3 we present results and demonstrate that, given an IBM representing an ideal experiment, we are able to identify IBM parameters for agent motility and adhesion. Following this, we replace our IBM representing an ideal experiment with an IBM that simulates a practically realisable experiment. In doing so we show that parameters cannot be successfully identified using ABC given the current experimental design. To improve parameter identifiablity we compare different experimental designs, and show that identification of IBM parameters is made more accurate if the size of the domain upon which the experiment is performed is expanded, as opposed to increasing the number of experimental repeats. Experimentally, expanding the size of the domain is equivalent to increasing the field of view of the microscope used to collect the experimental data. For instance, five simulation repeats on a larger domain provides more accurate identification of IBM parameters than 500 simulation repeats on a smaller domain. In Section 4 we discuss the results presented in this work.

## 2 Methods

In this section we first introduce the IBM. We then define what we mean by summary statistics and explain ABC and its implementation.

### 2.1 Individual-based model

An IBM is a computational model for simulating the behaviour of autonomous agents. The agents in the IBM represent cells, and each agent is able to move and interact with other agents. The IBM is simulated on a two-dimensional square lattice with lattice spacing Δ [18] and size *L_x_* by *L_y_*, where *L_x_* is the number of lattice sites in a row, and *L_y_* is the number of sites in a column. Each agent is initally assigned to a lattice site, from which it can move into adjacent sites. If an agent attempts to move into a site that is already occupied by another agent, the movement event is aborted. Processes such as this whereby one agent is allowed per site are often referred to as exclusion processes [18]. In the IBM time evolves continuously, in accordance with the Gillespie algorithm [19], such that agent movement events are modelled as exponentially distributed reaction events in a Markov chain. Attempted agent movement events occur with rate *P_m_* per unit time. *P_m_δt*, therefore, is the probability of an agent attempting to move in the next infinitesimally small time interval *δt*. A lattice site is denoted by *v* = (*i, j*), where *i* indicates the column number and *j* the row number. Each lattice site has four adjacent lattice sites (except for those sites situated on nonperiodic boundaries), and so the number of nearest neighbour lattice sites that are occupied by an agent, denoted by *n*, is 0 ≤ *n* ≤ 4. We denote the set of unoccupied nearest neighbour lattice sites by 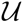.

The IBM domain size for simulations representing ideal experiments is *L_x_* = 100 by *L_y_* = 100, and the lattice sites indexed by 1 ≤ *j ≤ L_y_* and 1 ≤ *i* ≤ 10, and 1 ≤ *j* ≤ *L_y_* and 91 ≤ *i* ≤ *L_x_* are initially occupied by agents. In Fig. 1 the initial conditions in the IBM for the ideal experiment can be seen. The initial condition in Fig. 1 represents a ‘wound’, in that agents are positioned either side of a space, the ‘wound’, that they can migrate into. The agent migration into this space simulates one aspect of the wound-healing process. We refer to this simulation as ideal because the symmetry of the initial conditions may not be possible in a realistic experimental setting. The initial condition is also ideal as it is ‘double-sided’, as opposed to the ‘singlesided’ experiment data that we will later analyse. It has been shown that double-sided initial conditions can provide more information than single-sided initial conditions for some model parameters [11]. For instance, when increasing the number of agents in a simulation improves parameter identifiability.

**Figure 1:**
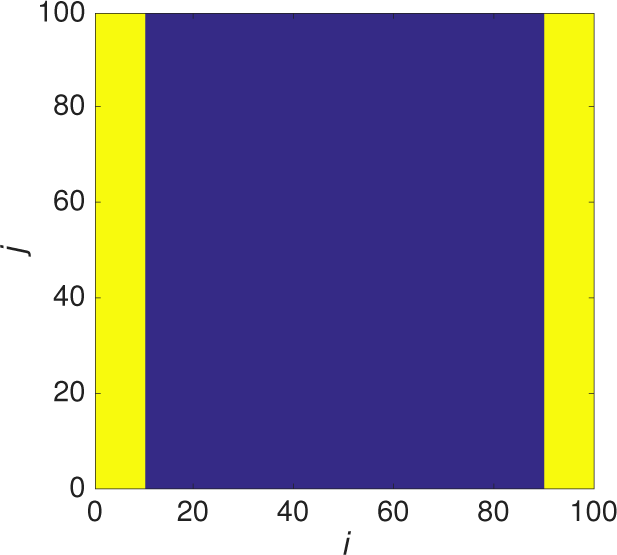
The initial condition in the IBM for the ideal experiment. Yellow indicates a site occupied by an agent and blue indicates an empty lattice site.

For the IBM of an ideal experiment all simulations have periodic boundary conditions at the top and bottom of the domain (i.e. for lattice sites indexed by *j* = 1 or *j* = *L_y_*), and no-flux boundary conditions at the left-hand and right-hand boundaries of the domain (i.e. for lattice sites indexed by *i* = 1 or *i* = *L_x_*).

### 2.2 Cell-cell adhesion models

In the IBM cell-cell interactions are simulated by altering the probability of an agent attempting to move, depending on the number of nearest occupied neighbours, *n*, an agent has. We employ two models to simulate cell-cell interactions in the IBM, one of which has been published before [20, 21]. We define *T*(*v′*|*v*) as the transition probability of an agent situated at site *v*, having been selected to move, attempting to move to site *v′*, where *v′* indicates one of the nearest neighbour sites of *v*. Therefore, *T*(*v′*|*v*) is only non-zero if *v* and *v′* are nearest neighbours. The transition probability in the first model, which we refer to as model A, is defined as

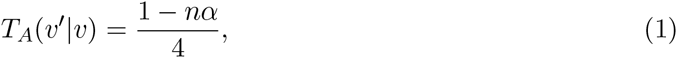

where *α* is the adhesion parameter. The subscript A on the transition probability in Eq. (1) indicates that this is the transition probability for model A. If *α* > 0 Eq. (1) models cell-cell adhesion, and if *α* < 0 Eq. (1) models cell-cell repulsion. The transition probabilities stated in Eq. (1) must satisfy

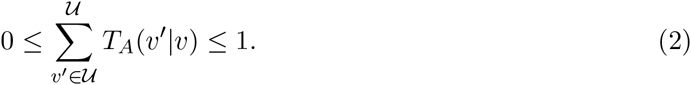

Equation (2) ensures the probability of an agent, if selected to move, attempting to move to any of its unoccupied nearest neighbour sites never exceeds unity, and so constrains the value *α* can take. The transition probability in the second model, which we refer to as model B [20, 21], is defined as

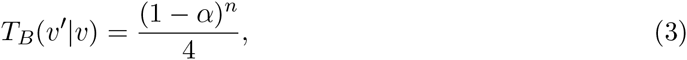

and must satisfy

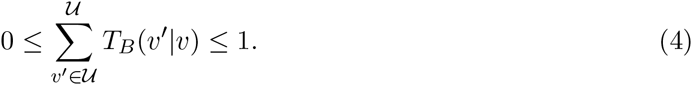

As in model A if *α* > 0 Eq. (3) models cell-cell adhesion, and if *α* < 0 Eq. (3) models cell-cell repulsion.

Models A and B simulate different forms of cell-cell interaction. In model A the transition probability is a linear function of *n*. Meanwhile, in model B the transition probability is a nonlinear function of *n*. Not only may these different forms of cell-cell interaction be relevant for different cell types, but implementing two models of cell-cell interaction allows us to test the robustness of the methods we present in this work.

### 2.3 Summary statistics

Summary statistics are lower-dimensional summaries of data that provide a tractable means to compare different sets of data. Summary statistics are important because experimental data is often of high dimensionality, and if we want to use experimental data to efficiently guide computational algorithms we require ways to accurately summarise it. We now define the summary statistics we apply to the IBM output and experimental data. Following this we describe how we utilise these summary statistics to implement ABC.

We initially use three summary statistics to evaluate the IBM output, all of which have been considered previously [9, 22]. The reason we study three summary statistics is to ascertain which summary statistic is most effective for the identification of agent motility and adhesion parameters in the IBM. These summary statistics are as follows:

#### Average horizontal displacement of agents

The average horizontal displacement of all agents, 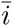, in a given time interval, [*t_i_, t_f_*], in the IBM is calculated as

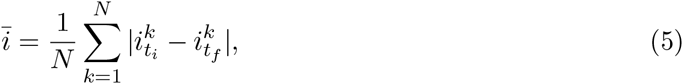

where 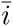 is the average horizontal displacement of agents, *N* is the total number of agents in the simulation, 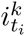 is the column position of agent *k* at time *t_i_* and 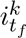 is the column position of agent *k* at time *t_f_*. We only look at the horizontal displacement of agents as this is the direction in which the majority of agent displacement will occur, due to the initial conditions of the IBM (Fig. 1). It has previously been shown that different cell-cell interactions have different effects on the average displacement of agents in an IBM [21]. As may be expected, repulsive (adhesive) interactions between agents tend to increase (decrease) the average displacement of agents, and so the average displacement of agents may be a useful summary statistic for distinguishing between repulsive and adhesive cell-cell interactions in the IBM.

#### Agent density profile

The agent density profile at time *t* in the IBM is calculated as:

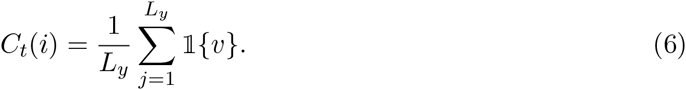

Here *C_t_*(*i*) is the agent density profile and 1 is the indicator function for the occupancy of a lattice site *v* (i.e. 1 if an agent occupies lattice site *v*, and 0 if it is not occupied by an agent). We have shown previously that different cell-cell interactions have different effects on the agent density profile [21]. For instance, repulsive interactions between agents can create a concave agent density profile, whereas adhesive interactions between agents can create a convex agent density profile. Therefore, the agent density profile may be an effective summary statistic for distinguishing between repulsive and adhesive cell-cell interactions in the IBM.

#### Pairwise-correlation function

The final summary statistic we consider is the pairwise-correlation function (PCF). The PCF provides a measure of the spatial clustering between agents in an IBM, and has been used frequently in the analysis of cell migratory processes [4, 9, 23, 24]. The PCF has also been successfully used as a summary statistic for the parameterisation of IBMs of cell migration [10]. We use 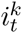 to denote the column position of agent *k* at time *t*, 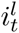 to denote the column position of agent *l* at time *t*, and define *c_t_*(*m*) to be the number of occupied pairs of lattice sites for each *nonperiodic*^4^ horizontal pair distance *m* = 1,…, *L_x_* – 1 at time *t*. This means *c_t_*(*m*) is given by

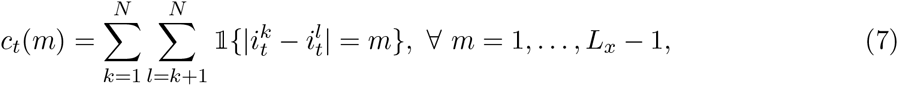

where 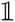 is the indicator function such that it is equal to 1 if 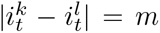, and is equal to 0 otherwise. In Eq. (7) only the pair agent distances in the horizontal direction are counted. Given the translational invariance of the initial conditions in the vertical direction of the IBM, the majority of important spatial information will be in the horizontal direction^5^. Binder and Simpson [24] demonstrated that is necessary to normalise Eq. (7) to account for volume exclusion. The normalisation term is

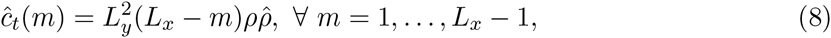

where *ρ* = *N*/(*L_x_L_y_*), 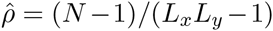. Equation (8) describes the expected number of pairs of occupied lattice sites, for each nonperiodic horizontal pair distance *m*, in an agent population distributed uniformly at random on the IBM domain. Combining Eqs. (7) and (8), the PCF is

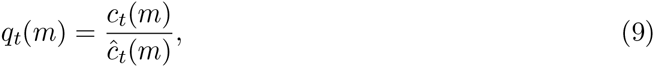

where *q_t_*(*m*), the PCF, is a measure of how far *c_t_*(*m*) departs from describing the expected number of occupied lattice pairs for each horizontal distance of an agent population spatially distributed uniformly at random on the IBM domain.

### 2.4 Approximate Bayesian computation

Here we introduce our ABC algorithm [14]. We define *M* as a stochastic model that takes parameters Θ and produces data *D*. This relationship can be written as *D* ~ *M*(Θ). For the IBM presented in this work Θ = (*P_m_, α*), where Θ is sampled from a prior distribution, *π*, and so this relationship can be written as Θ ~ *π*. The relationship between *π* and Θ is often written as Θ ~ *π*(Θ), which indicates that a new Θ sampled from the prior distribution may depend on the previous Θ. This relationship will be relevant later on in this work, however, initially each Θ sampled from *π* is independent of the previous Θ.

The identification of IBM parameters in this work centres around the following problem: given a stochastic model, *M*, and data, *D*, what is the probability density function that describes Θ being the model parameters that produced data *D*? More formally, we seek to obtain a posterior distribution, *p*(Θ|*D*), which is the conditional probability of Θ given *D* (and the model, *M*).

Typically, to compute the posterior distribution a likelihood function, *L*(*D*|Θ), is required. This is because the likelihood function and posterior distribution are related in the following manner by Bayes’ theorem:

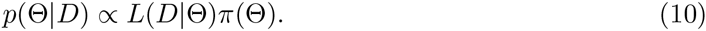

That is, the posterior distribution is proportional to the product of the likelihood function and the prior distribution. Approximate Bayesian computation is a well-known method for estimating posterior distributions of model parameters in scenarios where the likelihood function is *intractable* [14]. By an intractable likelihood function it is meant that the likelihood function is impossible or computationally prohibitive to obtain.

In many cases for ABC, due to the high dimensionality of the data, *D*, it is necessary to utilise a summary statistic, *S* = *S*(*D*). The summary statistics we employ in this work are of varying dimension. For instance, the agent density profile at time *t* has *L_x_* data points, whereas the average agent displacement at time *t* has one data point. Therefore we write *S*(*D*) as *S*(*D*)*_r,t_*, where *S*(*D*)*_r,t_* is the *r^th^* data point in the summary statistic at the *t^th^* sampling time.

The ABC method proceeds in the following manner: we wish to estimate a posterior distribution of Θ given *D*. We now simulate the process that created *D* using model *M* with parameters Θ, sampled from *π*, and produce data 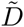. We calculate the difference between a summary statistic applied to *D* and 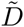 with

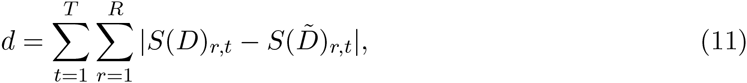

where *R* is the number of data points in *S*(*D*) and *T* is the number of sampling times. We repeat the above process many times, that is, sample Θ from *π*, produce 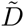 calculate *d* with Eq. (11), and only accept Θ for which *d* is below a user defined certain threshold (alternatively, a predefined number of Θ that minimise *d* can be accepted). This enables us to generate a distribution for Θ that is an approximation of the posterior distribution, *p*(Θ|*D*), given *M*. More specific details of the ABC algorithms we implement are introduced when necessary in the text.

## 3 Results

We begin by demonstrating that for an IBM representing an ideal experiment we are able to identify model parameters, given we use the appropriate summary statistics.

### 3.1 Ideal experiment

To ascertain the effectiveness of the chosen summary statistics to identify model parameters, we first attempt to identify Θ from data generated *synthetically*. Synthetic data is IBM data generated with fixed parameter values, and so can be thought of as a simulation equivalent of experimental data. To generate the synthetic data using the IBM we proceed as follows:

1. We choose parameters Θ to identify. To help clarify this explanation let us make these parameters Θ = (*P_m_,α*) = (0.5,0.1) in model A^6^.
2. For model A we perform a simulation of the IBM with Θ = (0.5,0.1), generate data, *D*, and calculate summary statistics, *S*(*D*), from the simulation at our time-points of interest. These times are *t* = [240,480, 720]. We choose these times as they are the times (in minutes) we will later analyse for the simulations of the practically realisable experiment, and correspond to 4 hours, 8 hours and 12 hours into an experiment.
3. We repeat step 2. ten times and calculate the ensemble average for each summary statistic for each individual time-point.

This procedure generates synthetic data for which we will now attempt to identify the parameters. In this work we present estimates for *P_m_* = 0.5 and *α* = 0.1 for model A, and *P_m_* = 0.5 and *α* = 0.25, and *P_m_* = 0.5 and *α* = –0.1 for model B. We examined identifying further combinations of values of *P_m_* and *α* from synthetic data. What we present here is a representative sample of the combinations we tested.

Throughout this work we sample *P_m_* and *α* for our model from uniform priors. In the case of model A, *P_m_* ∈ [0,1] and *α* ∈ [–0.2, 0.25], and for model B, *P_m_* ∈ [0,1] and *α* ∈ [−0.2,1.0]. We stipulate these lower and upper bounds for *α* for both models A and B to make sure inequalities (2) and (4) are satisfied.

We begin by implementing an ABC rejection algorithm since we expect to identify model parameters quickly as we are simulating an ideal experiment. The rejection ABC algorithm proceeds as follows:

1. Run 10^4^ IBM simulations, in each case using Θ sampled uniformly at random from the prior distributions.
2. Compute the distance *d* as defined in Eq. (11) for simulation times *t* = [240,480, 720].
3. Accept the 100 parameter values, Θ, that minimise *d*.

In Fig. 2 the posteriors generated using each of the three summary statistics applied to data from simulations of an ideal experiment are displayed. The most effective summary statistic for identifying the synthetic data parameters is the PCF. The effectiveness of the PCF for parameter identification is evident in the location of the posterior distribution density relative to the red dot (the red dot represents the synthetic data parameter values), and the narrow spread of the posterior distribution density as indicated by the scale bar in Fig. 2 (c), (f) and (i). The agent density profile summary statistic performs less well than the PCF for parameter identification, especially for model A (Fig. 2 (b)). In the case of the average agent displacement many combinations of *P_m_* and *α* lead to the same average agent displacement, which results in an extended region of possible parameter values. To some extent this is to be expected, as increasing either *P_m_* or *α* will have opposing effects on the average agent displacement. This means that using agent displacement as a summary statistic results in parameter identifiability issues in this example.

**Figure 2:**
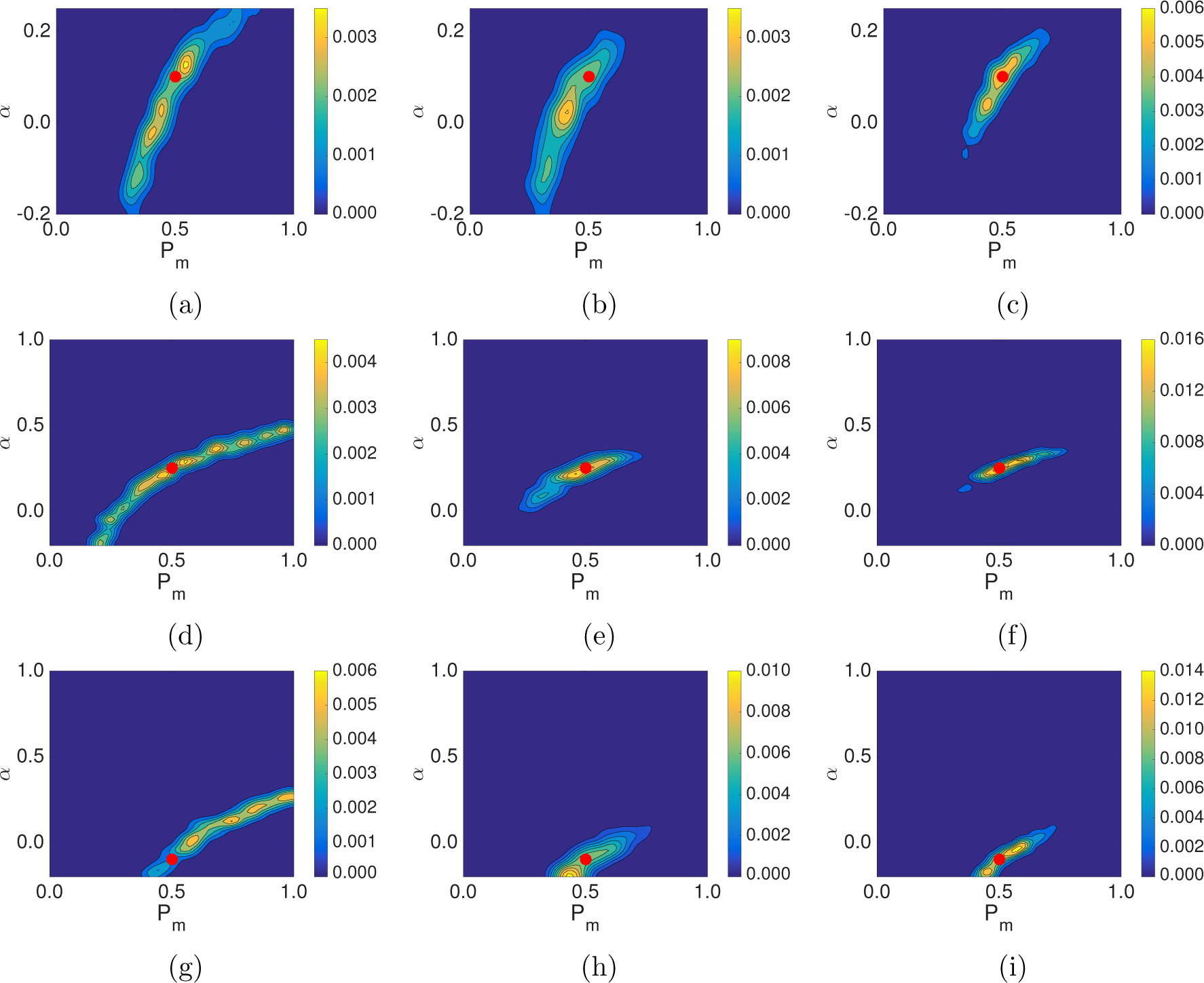
(a)-(c) Posterior distributions for model A for an ideal experiment with different summary statistics: (a) average displacement of agents in the horizontal direction; (b) agent density profile; (c) PCF. In all cases the red dot indicates the value of the parameters used to generate the synthetic data, *P_m_* = 0.5, *α* = 0.1. As indicated by the colour bar the yellow regions indicate areas of high relative density of the posterior distribution, while the blue regions indicate areas of low relative density of the posterior distribution. (d)-(f) Model B, *P_m_* = 0.5, *α* = 0.25: (d) average displacement of agents in the horizontal direction; (e) agent density profile; (f) PCF. (g)-(i) Model B, *P_m_* = 0.5, *α* = –0.1: (g) average displacement of agents in the horizontal direction; (h) agent density profile; (i) PCF.

To quantify the difference between the performance of the different summary statistics we use > the Kullback-Leibler divergence, which is a measure of the information gained in moving from ‘ the prior distribution to the posterior distribution [25]. The Kullback-Leibler divergence for a discrete probability distribution is defined as follows:

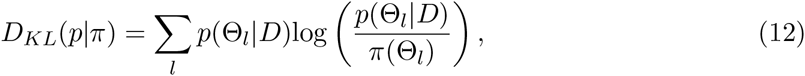

where the index *l* accounts for all possible discretised parameter pairs (i.e. all combinations of *P_m_* and *α*). A larger *D_KL_*(*p*|*π*) value suggests that more information is obtained (the entropy of the distribution is reduced) when moving from the prior distribution to the posterior distribution^7^. We discretise our posterior distribution onto a lattice with 2^6^ equally spaced values of *P_m_* and 2^6^ equally spaced values of *α*.

Computing *D_KL_*(*p*|*π*) for all nine plots in Fig. 2 gives: (a) 1.77; (b) 1.70; (c) 2.32; and (d) 2.15; (e) 2.57; (f) 3.35; and (g) 2.45; (h) 2.72; (i) 3.27. In tandem with the proximity of the peak of the posterior distribution densities to the red dots in Fig. 2 (c), (f) and (i), compared to Fig. 2 (a)-(b), (d)-(e) and (g)-(h), this suggests that the PCF summary statistic is more effective for parameter identification than the average agent displacement and agent density profile summary statistics.

### 3.2 Practically realisable experiment

In the previous section we demonstrated that for ideal experimental conditions the PCF summary statistic is best able to identify synthetic data parameters (for an IBM of an ideal experiment), and so moving forward we will only use the PCF summary statistic for parameter identification. Previous work has combined summary statistics to improve parameter identification [10]. However, in this case it makes a negligible improvement to the posterior (results not shown)^8^.

We now replace our IBM that represents an ideal experiment with an IBM that represents an actual experiment, and examine if synthetic data parameters can be identified in the IBM. We provide brief details of the experiment here, however, a more detailed description can be found in the supplementary material. In Fig. 3 a typical initial frame of the experimental data can be seen.

**Figure 3:**
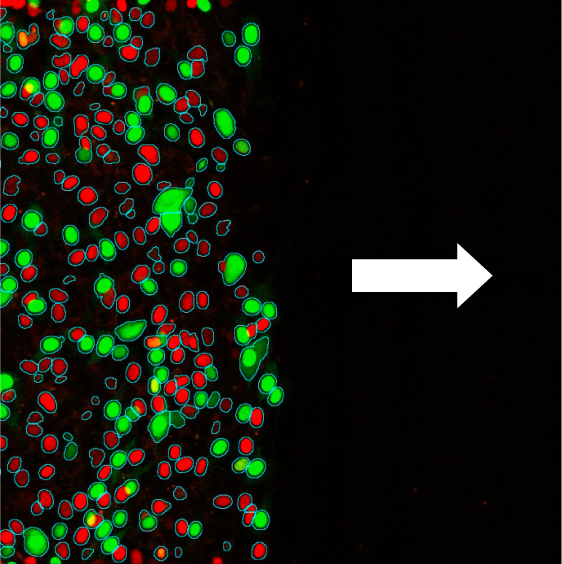
Typical initial frame of the experimental data. The cells are positioned such that they will migrate primarily horizontally into the space without cells, this space represents a wound (the direction of migration is indicated by the white arrow). The red and green cells are the same cells, with red and green indicating which phase of the cell cycle cells are in. In this work we do not take the cell cycle into account.

In total we have data from five repeats of the experiment. Each data set contains cell track data for every cell for sixty-four hours imaged at twenty minute intervals. Therefore, we have the information required to apply our summary statistics to the experimental data. More specifically, we have the position of all cells at each time interval so that the PCF may be computed.

One key difference between the ideal and practically realisable experiments is the size of the domain and, because of this, the number of agents in a simulation. As we have data from five experiments we now generate our synthetic data from five repeats of the IBM, using the same procedure as described in Section 3.1.

The experimental images were captured by a microscope with a field of view of 597.24 μm by 597.24 μm. The cell size in the experimental images is consistent with each cell occupying a 26 μm by 26 μm square lattice site. Given the size of the microscope field of view this means the IBM domain size is *L_x_* = 23 by *L_y_* = 23. We use the average initial conditions from the experiment to generate the initial conditions in the IBM. Exact details of how the initial condition is generated in the IBM, and how experimental data is mapped to a lattice can be found in the supplementary material.

We also alter the IBM to have flux (nonperiodic) boundary conditions at the left-hand and right-hand boundaries of the domain (i.e. for lattice sites with *j* = 1 or *j* = *N_y_*). The left-most column is kept at or above a constant density throughout the simulation time course. That is,after any movement event from the left-most column in the simulation the column density of the left-most column is calculated, and if found to be below a certain density agents are added to empty sites in this column chosen uniformly at random until the required density is achieved. This mechanism ensures that the agent density profile in the IBM replicates the evolution of the experimental data throughout the simulation. Further details regarding the implementation of this boundary condition are provided in the supplementary material. The top and bottom boundaries of the IBM domain remain periodic as cells were seen to move in and out of the microscope field at these boundaries in the experimental images, at an approximately equal rate.

To reduce the computational time of the ABC algorithm we now employ the Metropolis-Hastings algorithm. We do not implement rejection ABC as we expect parameter identification to be less efficient with a more realistic model, and so we implement a sequential Monte Carlo method. Given our model assumptions our implementation of the Metropolis-Hastings algorithm reduces to a Markov chain Monte Carlo method with a correlated outcome [14], of which we attempt 10^6^ realisations. Details of the implementation of the algorithm are given in the supplementary material. As before we sample from uniform priors *P_m_* ∈ [0,1] and *α* ∈ [−0.2, 0.25] for model A, and *P_m_* ∈ [0,1] and *α* ∈ [−0.2,1.0] for model B, and collect simulation data at *t* = [240, 480, 720]. We collect simulation data at three time-points so that the computational time is of practical length (our longest ABC implementations took approximately 192 hours). A value of *P_m_* = 0.5, given that the simulation time is in minutes, and the length of a lattice site is 26 μm, means that the motility of the agents is biologically realistic. To be precise, the agents here are approximately five times faster than cell motility rates previously published [4, 9]^9^. However, the cells considered in [4, 9] are not thought to exhibit cell-cell adhesion, and so a higher motility rate is sensible as agent movement is being reduced in the case of cell-cell adhesion in our IBM.

In Fig. 4 it can be seen that the synthetic data parameters cannot be accurately identified using our ABC method, with the PCF summary statistic, given the current IBM design. This is evident in the location of the red dots relative to the posterior distributions, and the wide spread of the posterior distributions as indicated by the scale bar in Fig. 4. A possible reason why the synthetic data parameters cannot be identified is that the synthetic data does not accurately represent the parameter values used to generate it, making parameter identification infeasible. To examine this possibility we calculated the variance in the PCF synthetic data^10^. In Fig. 5 (a)-(c) the blue line indicates the variance in the PCF synthetic data for the current simulation design generated from five repeats of the IBM on a domain of size *L_x_* = 23 by *L_y_* = 23.

**Figure 4:**
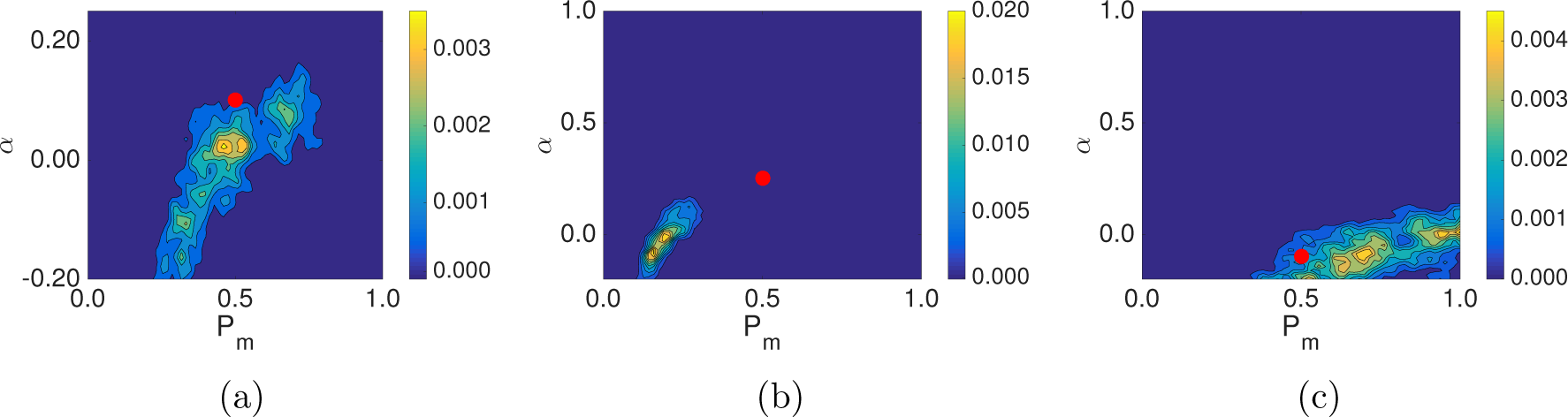
Posterior distributions for simulations of the experiment described in Section 2.5 using the PCF as a summary statistic for an IBM of size *L_x_* = 23 and *L_y_* = 23. The synthetic data is generated from five repeats of the IBM. (a) Model A: *P_m_* = 0.5, *α* = 0.1, (b) model B: *P_m_* = 0.5, *α* = 0.25, (c) model B: *P_m_* = 0.5, *α* = –0.1. In all cases the red dot indicates the value of the parameters used to generate synthetic data.

If the variance in the summary statistics of the synthetic data precludes accurate identification of model parameters using ABC, a sensible strategy may be to examine methods to reduce the variance in the summary statistics of the synthetic data. Reducing the variance of the summary statistics may mean the synthetic data is a more accurate reflection of the parameters values used to generate it. This may also explain why parameter identification for the ideal experiment was successful, as the variance in the summary statistics of the synthetic data was much smaller than for the practically realisable experiment (data not shown).

We conjectured that the variance in the summary statistics of the synthetic data could be reduced in two ways:

1. increasing the number of IBM repeats used to generate the synthetic data;
2. increasing the size of the IBM domain while keeping the column density of the initial conditions invariant. An example of this proposed initial condition is given in Fig. 6 (b). Importantly, increasing the size of the IBM domain increases the number of agents in the simulation, and can be thought of as equivalent to increasing the field of view of the microscope.

**Figure 5:**
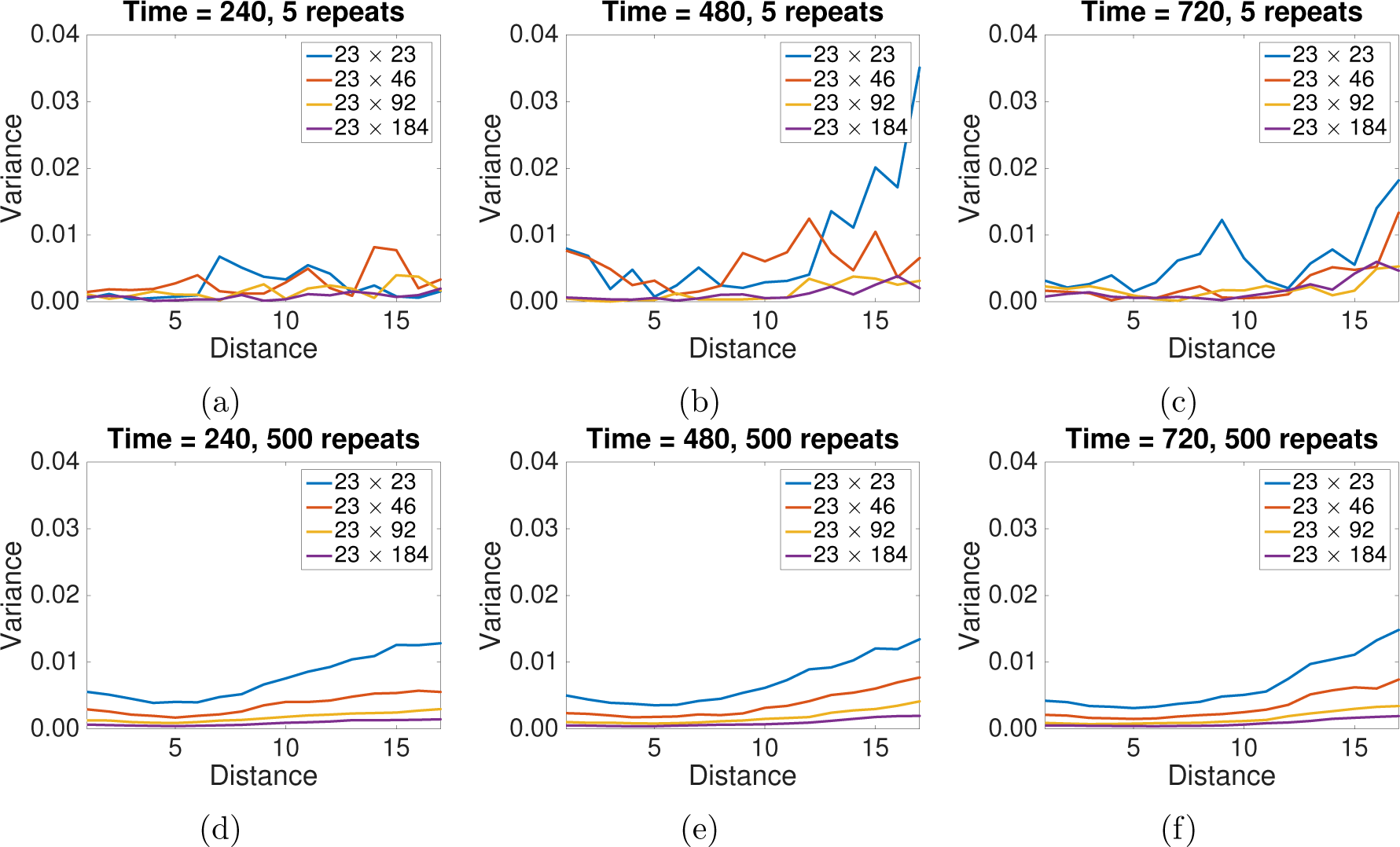
The variance in the PCF synthetic data for model B with *P_m_* = 0.5, *α* = 0.25 and different IBM domain sizes. Panels (a)-(c) display synthetic data generated from five repeats of the IBM, panels (d)-(f) display synthetic data generated from 500 repeats of the IBM. The domain size is indicated in the legend.

**Figure 6:**
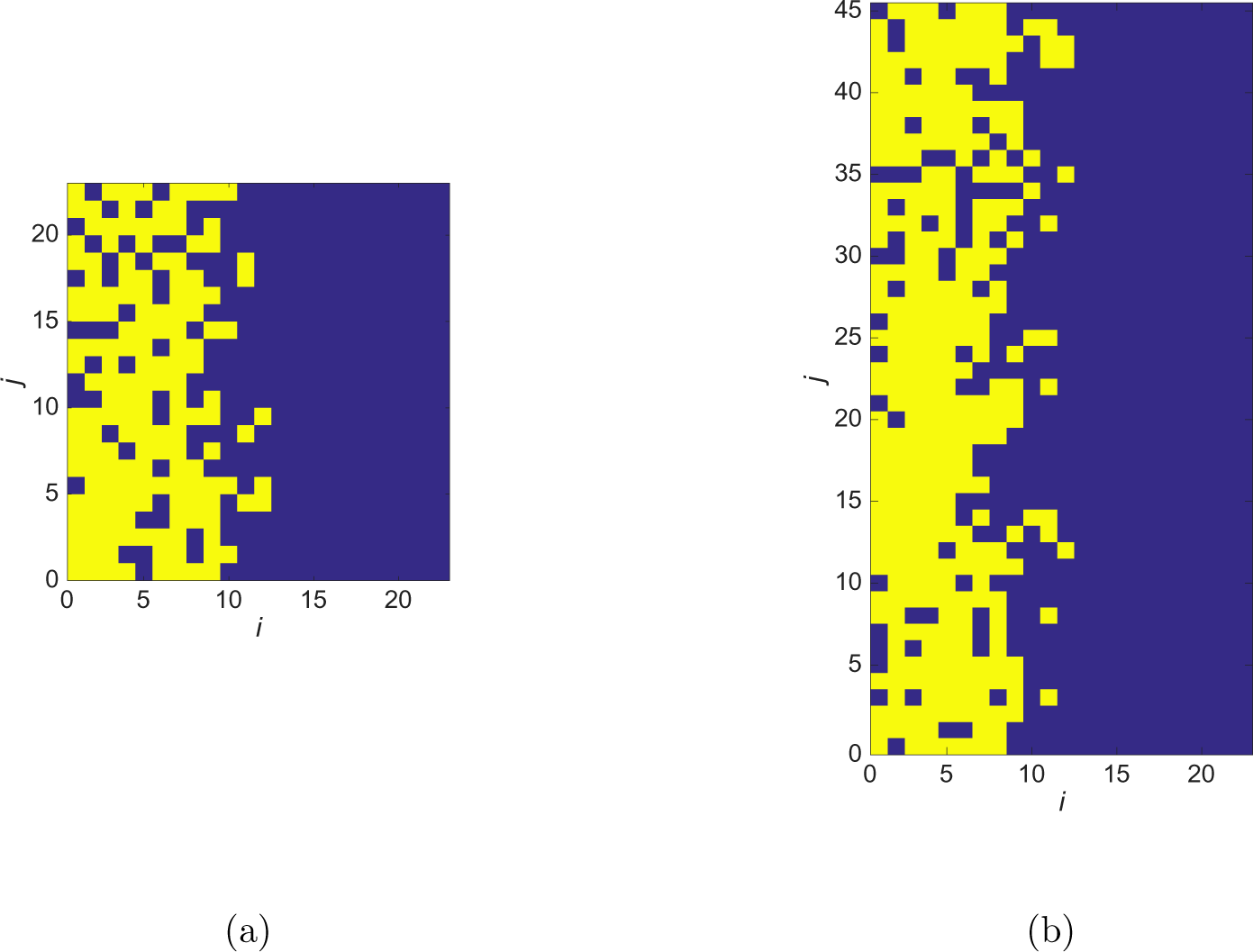
Increasing the size of the simulation domain while keeping the initial column densities the same. Panel (b) is twice the size of panel (a), however, the average initial density of each column is the same for both panels (a) and (b).

In Fig. 5 the variance in the PCF synthetic data for model B with *P_m_* = 0.5 and *α* = 0.25 for different domain sizes and varying numbers of repeats can be seen. It is evident that the variance in the PCF calculated from 500 repeats of a *L_x_* = 23 by *L_y_* = 23 sized domain (blue line in Fig. 5 (d)-(f)) is greater than the variance in the PCF calculated from five repeats of a *L_x_* = 23 by *L_y_* = 184 sized domain (purple line in Fig. 5 (a)-(c)). This can be understood by considering Eq. (7): the number of occupied lattice pairs for each horizontal pair distance used to generate the PCF does not increase linearly with the number of agents. Specifically, the number of occupied lattice pairs for each horizontal pair distance that generates the PCF is proportional to^11^

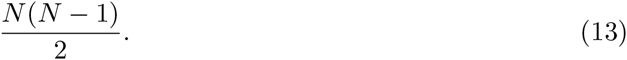

Therefore, the identification of parameters in experimental data using the PCF as a summary statistic may be best facilitated by increasing the size of the domain upon which the experiment is performed, rather than increasing the number of repeats of an experiment with a smaller domain. Further variance plots for models A and B for the PCF summary statistic can be found in the supplementary material.

It is important to note that it is also the case for the *agent density profile* synthetic data. If generated from 500 repeats of a *L_x_* = 23 by *L_y_* = 23 sized domain, the agent density profile synthetic data will have greater variance than the agent density profile synthetic data generated from five repeats of a *L_x_* = 23 by *L_y_* = 184 sized domain (data not shown). In this case it is an artefact of the lattice-based model. This is because the density of each column in the IBM can take on a greater range of values between 0 and 1 as the column length is increased, leading to a reduction in variance in the agent density profile synthetic data (especially in the initial conditions of the simulations used to generate the synthetic data). However, as we do not use the agent density profile summary statistic to identify parameters in the current simulation design we do not pursue this matter further.

### 3.3 Improving the experimental design

We now confirm that more accurate identification of synthetic data parameters occurs by expanding the domain upon which the experiment is performed, as opposed to increasing the number of experimental repeats.

In Fig. 7 (a)-(c) we plot the posterior distribution for synthetic data generated from 500 repeats of a *L_x_* = 23 by *L_y_* = 23 sized domain, while in Fig. 7 (d)-(f) we plot the posterior distribution generated from synthetic data generated five repeats of a *L_x_* = 23 by *L_y_* = 184 sized domain. As predicted, it is apparent that increasing the domain size is more effective for parameter identification than increasing the number of repeats used to generate the synthetic data. This is evident in the location (and narrow spread) of the posterior distribution relative to the red dot, whereby the posterior distribution is closer to the red dot in the case of Fig. 7 (d)-(f) compared to Fig. 7 (a)-(c). Despite this, the identification of the parameters for repulsive interactions remains somewhat elusive (Fig. 7 (f)). A possible reason for this is that the repulsive interaction we present here is a weak one, due to the constraint of Eqs. (2) and (4), and larger values of |*α*| are easier to identify as they have a more profound effect on the behaviour of the agent population.

**Figure 7:**
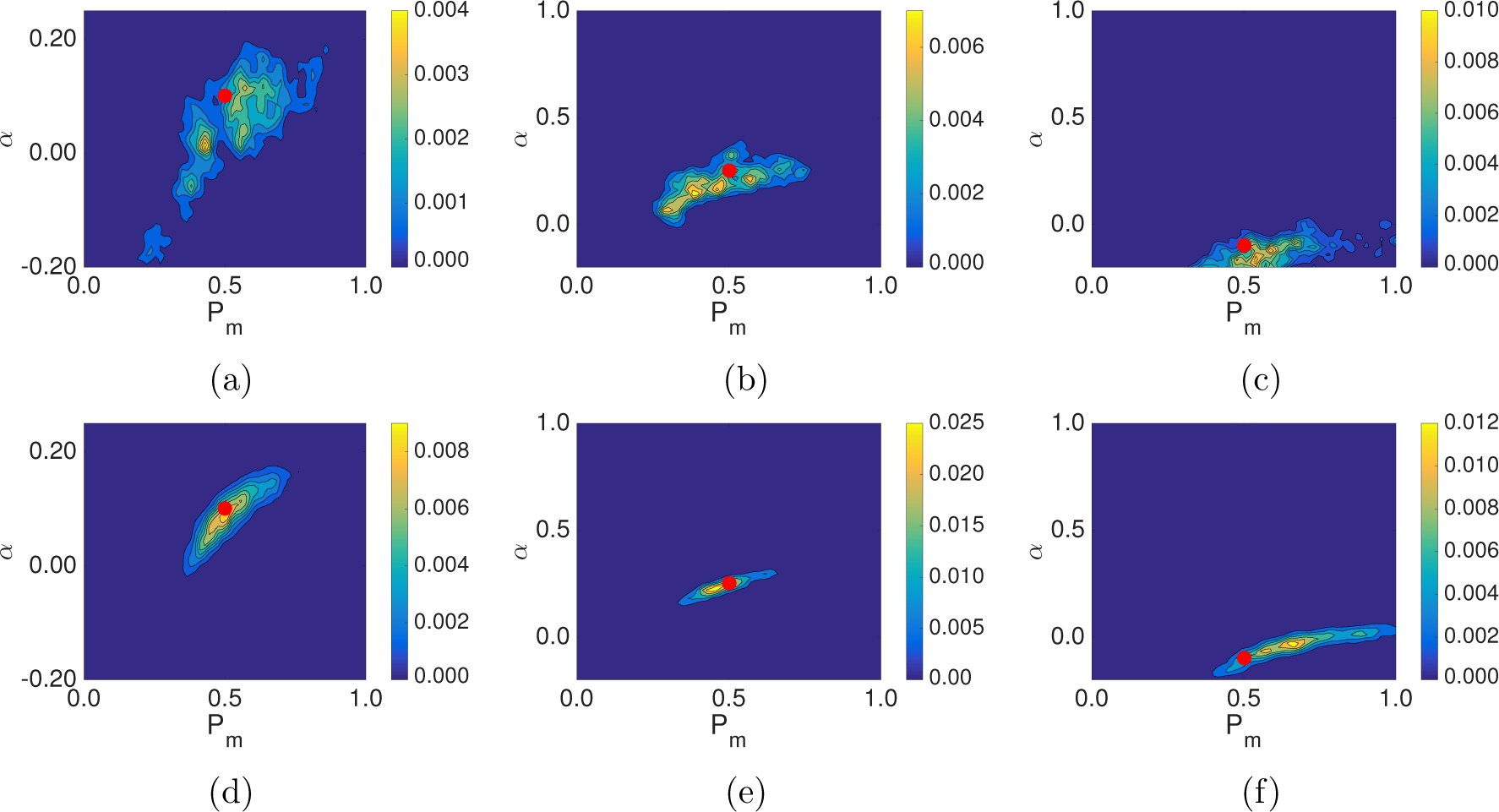
(a)-(c) Posterior distributions for simulations of the experiment using the PCF as a summary statistic for an IBM simulated on a domain of size *L_x_* = 23 by *L_y_* = 23 with synthetic data generated from 500 repeats. (a) Model A: *P_m_* = 0.5, *α* = 0.1, (b) model B: *P_m_* = 0.5, *α* = 0.25, (c) model B: *P_m_* = 0.5, *α* = –0.1. (d)-(f) Posterior distribution plots for simulations of the experiment using the PCF as a summary statistic for an IBM simulated on a domain of size *L_x_* = 23 by *L_y_* = 184 with synthetic data generated from five repeats. (a) Model A: *P_m_* = 0.5, *α* = 0.1, (b) model B: *P_m_* = 0.5, *α* = 0.25, (c) model B: *P_m_* = 0.5, *α* = –0.1. Further figure information can be found in Fig. 4.

Computing *D_KL_*(*p*|*π*) for all six plots in Fig. 7 gives: (a) 2.55; (b) 2.69; (c) 1.53; and (d) 3.69; (e) 2.97; (f) 3.54. In tandem with the proximity of the peak of the posterior distribution densities to the red dots in Fig. 7 (d)-(f) compared to Fig. 7 (a)-(c), this suggests that generating synthetic data on a larger domain is more effective for improving parameter identification than increasing the number of repeats used to generate the synthetic data.

## 4 Discussion

In this work we have presented methods to identify motility and adhesion parameters in an IBM of a wound-healing assay. Our findings suggest that for a commonly performed experiment increasing the size of the experimental domain can be more effective in improving the accuracy of parameter identification, when compared to increasing the number of repeats of the experiment. This is because increasing the size of the domain, which is equivalent to increasing the number of cells in the experiment, more effectively reduces the variance in the synthetic data from which the parameters are identified. The reason for this reduction in variance is explained by Eq. (7), where the number of agent pair counts that generate the PCF increases nonlinearly with the number of agents on the domain. In addition, increasing the size of the experimental domain may make the collection of experimental data less time-consuming, as potentially fewer repeats of the experiment will have to be conducted. For instance, five repeats of the experiment on a larger domain provides more information about parameters than 500 repeats of the experiment on a smaller domain (in the examples we have presented in this work).

We also studied using the average horizontal displacement of agents and the agent density profile as summary statistics. These were found to be less effective than the PCF in parameter identification. This was especially the case for the averaged agent displacement, whereby a range of adhesion and motility parameters could result in the same average agent displacement. This result suggests that agent displacement may not be a suitable summary statistic for estimating cell motility and adhesion parameters, due to parameter identifiability issues.

The obvious extension to the work presented here is to experimentally validate the findings. That is, expand the wound-healing experimental domain and demonstrate: i) the cell migratory process can be effectively described by the model we have presented here; and ii) the experimental parameters are identifiable with a larger experimental domain. If validated, alterations could be made to the IBM to try and further improve parameter identification, and evidence may be provided that demonstrates which adhesion model, A or B, is more applicable to the cell type under consideration.

To conclude, the findings presented in this work will be of particular interest to those concerned with performing experiments that enable the effective parameterisation of cell migratory processes. In particular, cell migratory processes in which cell-cell adhesion or repulsion are known to play an important role. More generally, we have also suggested time and cost-saving alterations to a commonly performed experiment for identifying cell motility parameters.

## Acknowledgements

RJHR would like to thank the UK’s Engineering and Physical Sciences Research Council (EPSRC, EP/G03706X/1) for funding through a studentship at the Systems Biology programme of The University of Oxford’s Doctoral Training Centre. RLM was supported by a Medical Research Scotland Project Grant (436FRG). The authors declare no competing interests.

## Contributions

RJHR, REB and CAY conceived the work, and performed the mathematical and computational analysis. Data collection and analysis was performed by RLM and MJF. RJHR, REB and CAY drafted the manuscript. All authors agree with manuscript results and conclusions. All authors approved the final version.

## Footnotes

1 Wound-healing assays are also often referred to as scratch assays.

2 By an effective representation we mean the IBM captures the salient features of the process of interest, and is therefore a viable research tool with which to study the process of interest

3 Throughout this work we assume that cellular processes such as migration have constant parameter values associated with them.

4 By nonperiodic it is meant the distance measured between two agents cannot cross the IBM boundary.

5 This approach is in agreement with previous studies [24], which showed the most relevant information from the PCF summary statistic is perpendicular to the wound axis in a wound-healing assay.

6 A value of *P_m_* = 0.5, given that the simulation time will later be defined to be in minutes, and the length of a lattice site represents cell length (typically between 10μm-100μm), means that the motility of the agents is biologically realistic. The parameter *α* is dimensionless. The experimental realism of these parameters will be expanded on when we address the simulation of a practically realisable experiment.

7 However, this does not necessarily mean the posterior distribution is a more accurate representation of the parameter distribution.

8 That there is little improvement in parameter identification from combining summary statistics is to be expected. Combining summary statistics is most effective when the posterior distributions are ‘orthogonal’,which is not the case for the posterior distributions created by the summary statistics presented here.

9 ^9^Using the relationship that the diffusion coefficient is equal to *P_m_*Δ^2^.

10 The variance was calculated using MATLAB’s in-built var function.

11 This is not quite correct as a distance of ‘0’ between agents, that is they share the same column, is not accounted for in Eq. (7). To make Eq. (13) exact is not trivial as the expected number of agents each agent shares a column with depends on both the column position and simulation time.

